# Hydrogen bonds meet self-attention: all you need for general-purpose protein structure embedding

**DOI:** 10.1101/2021.01.31.428935

**Authors:** Cheng Chen, Yuguo Zha, Daming Zhu, Kang Ning, Xuefeng Cui

**Author notes:** Who contributed equally to this work (Cheng Chen, Yuguo Zha).

## Abstract

General-purpose protein structure embedding can be used for many important protein biology tasks, such as protein design, drug design and binding affinity prediction. Recent researches have shown that attention-based encoder layers are more suitable to learn high-level features. Based on this key observation, we treat low-level representation learning and high-level representation learning separately, and propose a two-level general-purpose protein structure embedding neural network, called ContactLib-ATT. On the local embedding level, a simple yet meaningful hydrogen-bond representation is learned. On the global embedding level, attention-based encoder layers are employed for global representation learning. In our experiments, ContactLib-ATT achieves a SCOP superfamily classification accuracy of 82.4% (i.e., 6.7% higher than state-of-the-art method) on the SCOP40 2.07 dataset. Moreover, ContactLib-ATT is demonstrated to successfully simulate a structure-based search engine for remote homologous proteins, and our top-10 candidate list contains at least one remote homolog with a probability of 91.9%. Source codes: https://github.com/xfcui/contactlib.

## I. Introduction

Proteins play critical functions in living organisms. In order to understand how proteins function, homologous proteins can be analyzed to find correlations between the conserved function and the conserved structure. This is the main reason that the SCOP database [1] is built and manually curated to hierarchically classify proteins. Specifically, close homologs sharing similar sequences are grouped as families, and remote homologs sharing similar structures (or functions) are grouped as superfamilies. Given an experimentally determined new protein structure, accurate and fast superfamily classification is a critical step for many biological studies [2], [3].

In the past two decades, many SCOP superfamily classification methods have been proposed. These methods can be divided into two categories: sequence-based and structure-based. For sequence-based methods [2], [4], [5], hidden Markov models (HMMs) are first built to represent superfamilies, and pairwise alignments [6], [7] between the query protein and the representative HMMs are then used to identify the nearest superfamily. For structure-based methods [8], [9], pairwise sequence alignments and pairwise structure alignments [10]–[12] between the query protein and each protein of a non-redundant SCOP database are first conducted, and the alignment similarities are then analyzed to classify the query protein. It can be seen that all these methods are based on database scanning to calculate pairwise similarities between the query protein and all SCOP superfamilies (represented as HMMs, sequences or structures).

The SCOP superfamily classification problem remains challenging as a significant portion of PDB remains unclassified [1]. As of January 28, 2021, 102, 550 PDB entries have been classified in SCOP 2.07-2021-01-09 [1], while 174, 014 PDB entries have been deposited in PDB [13]. This happens because SCOP employs a sequence-based homology search algorithm to automatically classify new proteins [1]. Consequently, new proteins that do not have close homologs in SCOP are difficult to be classified because remote homologs do not share similar sequences. Instead, protein structures are more reliable evidences to find remote homologs, but the pairwise structure alignment problem is proved to be NP-hard [14]. Although many heuristic algorithms have been implemented [10]–[12], database scanning with these heuristic algorithms is still not practical for timely tasks. Therefore, a structure-based SCOP superfamily classification algorithm without database scanning is needed for timely tasks.

Recent developments in deep neural networks (DNNs) enable new approaches to protein classifications and related protein bioinformatics tasks. For example, protein sequence embedding DNNs [15], [16] have been introduced for general-purpose, and protein structure embedding DNNs [17], [18] have been designed for alignment-free homology search. Moreover, Transformer DNNs [19] have been proposed for protein structure prediction [20], [21], protein design [22], drug design [23] and antigen-antibody binding prediction [24]. Note that a general-purpose protein structure embedding based on a Transformer DNN is still missing. Here, we would like to explore this highly promising approach, and demonstrate its advantages for structure-based SCOP superfamily classifications.

In this manuscript, we introduce a novel attention-based DNN, called ContactLib-ATT, to embed protein structures. As our initial study, we applied ContactLib-ATT for the SCOP superfamily classification problem [1]. More applications will be explored in the future. The new ContactLib-ATT has several key innovations comparing to previous methods in protein bioinformatics: (a) ContactLib-ATT employs a two level embedding method that incoperates a local embedding for local contact contexts and a global embedding for the global network of local contact contexts; (b) our local embedding is a general framework that accepts either sequential or pairwise features according to your definition of a contact context; (c) our global embedding is based on attention-based encoder layers [19] that has been shown to be a better choice to learn high-level features [25]; (d) our classification method directly classify protein structures without any database scanning and consequently the running time of ContactLib-ATT is less than one second.

In our experiments on SCOP 2.07 [1], ContactLib-ATT is compared to state-of-the-art DNN methods. Comparing to the convolution DNN introduced by DeepFold [17], ContactLib-ATT boost the superfamily classification accuracy from 75.7% to 82.4%. Especially, for the most challenging cases (i.e., short proteins with less than 64 residues), the superfamily classification accuracy is increased by 10.7%. All these results suggest that hydrogen bonds with self-attention are all you need for general-purpose protein structure embedding.

## II. Methods

The main idea of our ContactLib-ATT method is as following. A protein structure can be loosely defined as a network of amino acids connected by peptide bonds and hydrogen bonds. Here, peptide bonds form a chain structure that is common among all proteins. However, hydrogen bonds form a more complicated network structure that mimics secondary structures and the global topologies of secondary structure elements. Based on this key observation, local (i.e., close in 3D space) fragments around residue-residue contacts (including hydrogen bonds) have been successfully used as local fingerprints by alignment-free methods to find homologous proteins [17], [26]–[28]. The idea of ContactLib-ATT is one step further to combine these local fingerprints to a global one using an attention-based encoder neural network [19].

As shown in Figure 1a, a new ContactLib-ATT method is introduced in three steps. First, given a query protein structure, each hydrogen bond (H-bond) and its context is abstracted and converted into a local embedding vector (see Section II-A). Then, the query protein structure as an H-bond network is converted into a global embedding vector (see Section II-B). This is done by incorporating attention-based encoder layers [19]. Finally, the global embedding vector is used to classify the query protein structure into its SCOP superfamily (see Section II-C, [1]). Certainly, the global embedding vector can be easily adopted for other structure-based protein bioinformatics tasks, such as the homology search problem and the function annotation problem.

**Fig. 1.**
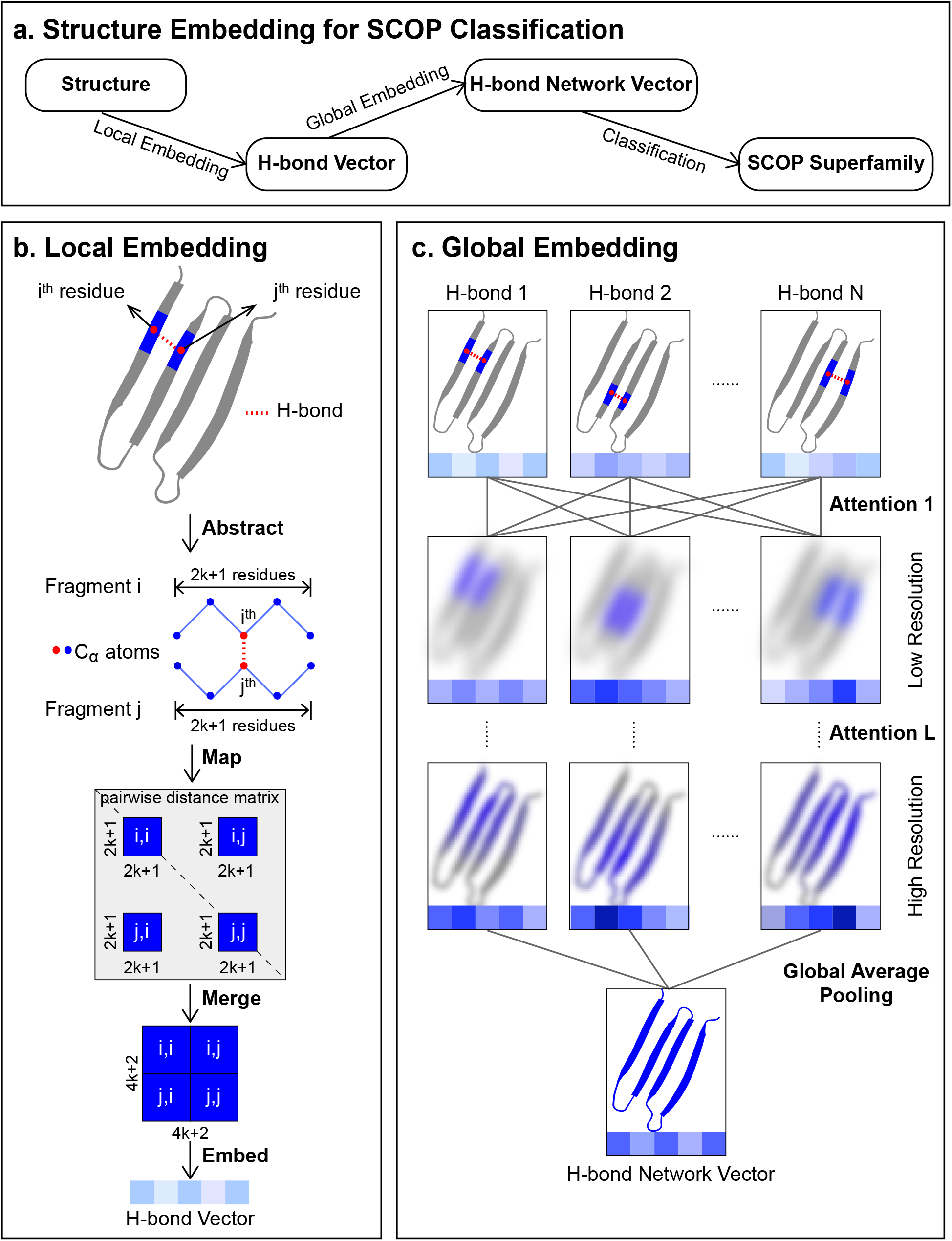
Illustrations of protein structure embedding and classification with ContactLib-ATT: (a) the pipeline of ContactLib-ATT has three steps; (b) the first local embedding step to abstract and to embed local hydrogen bond (H-bond) contexts is described in Section II-A; (c) the second global embedding step to embed a global H-bond network is described in Section II-B; and the third classification step (not shown in the Figure) as the first application of our general-purpose protein structure embedding is described in Section II-C.

The novel ContactLib-ATT model has at least two major advantages over state-of-the-art convolution neural network models, such as DeepFold [17]. First, the local patterns are learned by dense layers instead of convolution layers. By doing this, ContactLib-ATT is able to flexibly adopt more biologically meaningful local information, such as the context of an H-bond. Second, the global patterns are learned by attention-based encoder layers [19] instead of convolution layers. Recent researches have already shown that attention-based layers are more suitable to learn high-level (e.g., global and topological) features than convolution layers [25]. Indeed, DeepFold cannot learn contact patterns between residues that are far from the backbone chain (e.g., patch (*i, j*) of the distance matrix shown in Figure 1b) because such remote contacts cannot be covered by a small number of convolution layers. Introducing more convolution layers would also involve more padding (i.e., noises), which might become the majority of input signals for short proteins.

### A. Local embedding

The task of local embedding (as shown in Figure 1b) is to abstract the local context of any H-bond of a query protein structure, and then apply a deep neural network (DNN) to convert any H-bond context to its embedding vector. The H-bond context abstraction works as following. Initially, all H-bonds within a query protein structure are computed by DSSP. For the sake of simple explanations, we focus on processing the hydrogen bond between donor residue *i* and acceptor residue *j* (where *i* and *j* are residue indices defined by PDB, [13]). For ContactLib-ATT, the context of this H-bond is defined to be the *k* neighbor residues on either side of residue *i* or *j* on the backbone chain, and the *C*_*α*_ atoms are used as representatives of these 4*k* + 2 neighbor residues. Then, this H-bond context is mapped to four patches (i.e., sub-matrices) of the pairwise distance matrix between all *C*_*α*_ atoms, and the four patches are merged as one pairwise distance matrix to structurally represent the H-bond context. Previous researches have already shown that similar patches around residue-residue contacts carry critical fingerprint information to distinguish protein structures [17], [26]–[28], and ContactLib-ATT is based on this key observation.

Given an H-bond context, a DNN is applied to embed the pairwise distances between 4*k* + 2 residues (i.e., representative *C*_*α*_ atoms) to an H-bond vector. It can be shown that the pairwise distance matrix is symmetric, and hence only the upper triangle of the matrix is used as the input of DNN. Similar to DeepFold [17] and TMscore [29], [30], distance *d*_*a,b*_ between residues *a* and *b* is first converted to 1/(1 + (*d*_*a,b*_*/d*_0_)^*p*^), where *d*_0_ = 3.8 and *p* ∈ {1, 2, 3}. By doing this, relatively shorter distances have greater impacts to the embedded vector, and the impacts are upper bounded. Then, two dense blocks are employed for embedding, where each dense block contains a dense layer, a layer normalization [31] and a ReLU activation [32]. Here, the number of the output neurons equals to the dimension of the embedded H-bond vector (*E*), and the number of hidden neurons equals to 4*E*. Again, the main contribution of local embedding is introducing H-bond contexts instead of finding the best embedding DNN.

### B. Global embedding

Using our local embedding, *N* H-bond contexts of the query protein structure are embedded into *N* H-bond vectors. As shown in Figure 1c, the task of global embedding is to combine local H-bond vectors into a global H-bond network vector by an attention-based encoder neural network. Specifically, *L* attention-based encoder layers (i.e., Transformer encoder layers, [19]) are employed to embed *N* local H-bond vectors to *N* global H-bond network vectors, and a global average pooling layer [33] is adopted to reduce the *N* H-bond network vectors into a single one. To reduce the number of hyper-parameters of ContactLib-ATT, the dimension of the global embedding vector is set to be identical to that of the local embedding vector (*E*). Moreover, for all attention-based encoder layers, the number of heads is set to be *E*/64, and the dimension of the feed-forward network is set to be 4*E*. These settings are consistent with the original Transformer when *E* = 512 [19]. Our attention-based encoder neural network is based on the understanding of protein folding such that all H-bonds should contribute together to fold the protein into a compact and stable structure. Thus, these H-bonds should be more or less correlated, and an H-bond network can be virtually constructed to model such correlations (i.e., edges) between H-bonds (i.e., nodes). Here, the H-bond network is assumed to be a complete graph, and it is open to introduce more efficient or more effective sparse graphs [34], [35] to replace the complete graph. Based on the H-bond network, the attention-based encoder network is trained to understand the correlations between H-bonds, and to incorporate local H-bond embedding vector into global H-bond network embedding vector. As a result, the H-bond network is converted to an embedding vector as a representation of the query protein structure.

### C. SCOP superfamily classification with data augmentation and multi-tasking

Once the query protein structure is converted to an embedding vector, it can be easily used for many structure-based applications [17], [18], [36]. Here, one application to classify SCOP superfamilies [1] is demonstrated. Specifically, a dense layer and a softmax activation is simply used as our classification DNN. Given query protein structure *q*, let *y*_*q*_ be the true SCOP superfamily, *ŷ*_*q*_ be the predicted SCOP superfamily, and *v*_*q*_ be the embedded H-bond network vector. Then, the loss function for *q* is set to be *CE*(*y*_*q*_, *ŷ* _*q*_) + 0.1 × *RMS*(*v*_*q*_), where *CE* is the cross entropy loss and *RMS* is the root mean square loss. Here, the cross entropy loss has been widely used with softmax activation, and the root mean square loss has been used by DeepFold [17] as an embedding regularization. Now, it can be seen that ContactLib-ATT is an end-to-end deep learning model, and the embedded vectors are optimized for the classification task.

In order to maximize the utilities of the limited data, data augmentation techniques can be adopted. Here, it is important to understand the risk of using all available data: the trained model will be overfitted to query protein structures that have many close homologs in SCOP, which tend to be efficiently found by sequence alignments. Thus, it is safer to train the model with only remote homologs, but that would significantly reduce the size of training data. Current implementation of ContactLib-ATT incorporate two simple data augmentations. First, protein structures are pre-clustered by sequence similarities, and for each training epoch, only one structure is randomly selected from each cluster. Consequently, the model does not see close homologs frequently. Second, Gaussian noises are added to all atom coordinates of the selected protein structures so that even if the model sees the same protein structure multiple times, the actual input structures are slightly different.

In order to avoid overfitting, multi-tasking techniques can be adopted. Current implementation of ContactLib-ATT incorporates simple multi-tasking classifications on the SCOP class level, the SCOP fold level and the SCOP superfamily level at the same time. This approach is chosen because superfamily annotations implies class and fold annotations, and high-quality embedding for superfamily classification should also be good at class and fold classifications. Moreover, it provides possibilities for the ContactLib-ATT model to learn the underlying logic of the hierarchical SCOP classifications. In the future, we would like to evaluate more multi-tasking approaches for ContactLib-ATT.

## III. Results

To evaluate the classification accuracies of different methods, a subset of SCOP 2.07 [1] is built as following. First, protein domains of SCOP 2.07 are filtered by a maximum sequence identity of 40%, a minimum sequence length of 20, and a minimum H-bond count of 20. Then, protein domains from SCOP 2.06 (i.e., a subset of SCOP 2.07) are randomly divided into training and validation subsets. For each superfamily, one domain is randomly selected and reserved for training. For the unreserved domains, 20% are randomly selected as the validation dataset, and the remaining 80% are combined with the reserved domains as the training dataset. Finally, protein domains present in SCOP 2.07 but not in SCOP 2.06 are used as the testing dataset. As a result, 11, 029, 2, 280 and 272 protein domains are selected for training, validation and testing, respectively.

To demonstrate the advantages of the novel ContactLib-ATT model, three more deep learning methods are tested. For the first model, six convolution blocks and one global average pooling layer is adopted from DeepFold [17] to embed the query protein structure. Then, unlike DeepFold, the classification model introduced in Section II-C is adopted to classify the embedded structure. For the second model, the last two convolution blocks of the first model is replaced by two attention-based encoder layers [19]. For the last model, the attention-based encoder layers of ContactLib-ATT are replaced by the local embedding model introduced in Section II-A. These three models are simply referred as DeepFold, DeepFold-ATT and ContactLib-DNN in this study, and they are compared to our ContactLib-ATT in Table I.

**TABLE I.** Comparison of methodologies

|  | Input Features | Local Embedding | Global Embedding |
| --- | --- | --- | --- |
| DeepFold | Full Structure | Convolution | Convolution |
| DeepFold-ATT | Full Structure | Convolution | **Attention** |
| ContactLib-DNN | **H-bond Contexts** | DNN | DNN |
| ContactLib-ATT | **H-bond Contexts** | DNN | **Attention** |

In order to evaluate the contributions of different components described in Section II-C, four variants of ContactLib-ATT are tested. Specifically, ContactLib-ATT00 is the base model without data augmentation (DA) and multi-tasking (MT); ContactLib-ATT01 is the advance model with only MT; ContactLib-ATT10 is the advance model with only DA; and ContactLib-ATT11 (or simply ContactLib-ATT) is the complete model with both DA and MT. Moreover, our H-bond context includes *k* = 8 neighbor residues, our global embedding employs *L* = 3 attention-based encoder layers [19], and both local and global embedded vectors have a dimension of *E* = 1, 024. Increasing or decreasing one of these three hyper-parameters by a factor of two has no significant impacts on accuracies, and thus the results are not included here.

Finally, the experiments are designed as following. To make it fair, all tested deep learning models employ the same configuration of layer normalizations [31], dropout regularizations [37], ReLU activations [32], and classification models (described in Section II-C). Each method is first trained on the training dataset, and then evaluated on the validation dataset and the testing dataset. Since the tested methods can only predict SCOP superfamilies [1], the SCOP fold (or class) containing the predicted superfamily is presumed to be the predicted fold (or class). The accuracy of a model is defined as the percentage of the correctly predicted SCOP classifications (e.g., classes, folds or superfamilies) over all predictions. The above process is repeated five times and the mean accuracies and the standard deviations are calculated.

### A. Comparison to state-of-the-art methods

In this experiment, our ContactLib-ATT is compared to state-of-the-art deep learning methods. It is shown that ContactLib-ATT achieves the highest accuracies on both the validation dataset and the testing dataset. Moreover, the outstanding performance is mainly because of the H-bond contexts, the attention-based global embedding, the data augmentation and the multi-tasking introduced by ContactLib-ATT.

From Table II, it can be seen that ContactLib-ATT11 is always the most accurate method for all SCOP classifications on the validation dataset. For example, the mean superfamily classification accuracy of ContactLib-ATT11 is 82.4%, which is 6.7% higher than that of DeepFold [17]. Considering that the standard deviation of the accuracy is 0.5%, the improvement of 6.7% is statistically significant. Actually, without data augmentation and multi-tasking, ContactLib-ATT00 already out-performs other tested methods by at least 1.5% on superfamily classifications. Using either data augmentation or multi-tasking slightly improves the accuracies by up-to 0.8%. However, using both of them boosts the accuracies by 2.9%. Therefore, all of the deep learning model, the data augmentation and the multi-tasking of ContactLib-ATT contributes to the accuracy improvements.

**TABLE II.** Accuracies on the validation dataset

|  | Class | Fold | Superfamily |
| --- | --- | --- | --- |
| DeepFold | 91.5±0.3% | 79.8±0.2% | 75.7±0.3% |
| DeepFold-ATT | 92.8±0.2% | 81.5±0.4% | 78.0±0.4% |
| ContactLib-DNN | 92.0±0.5% | 78.5±0.5% | 76.0±0.4% |
| ContactLib-ATT00 | 93.6±0.3% | 81.8±0.5% | 79.5±0.3% |
| ContactLib-ATT01 | 94.5±0.3% | 82.9±0.1% | 80.2±0.3% |
| ContactLib-ATT10 | 93.6±0.3% | 82.6±0.4% | 80.3±0.5% |
| ContactLib-ATT11 | **94.5±0.2%** | **85.0±0.3%** | **82.4±0.5%** |

Using novel H-bond contexts instead of full structures is one reason why our ContactLib-ATT model outperforms existing methods. This is well illustrated by comparing the results of DeepFold-ATT and ContactLib-ATT00 in Table II. It can be seen that using H-bond contexts with the DNN-based local embedding (introduced in Section II-A) produces more accurate predictions than using full structures with the convolution local embedding (introduced by DeepFold, [17]). One possible explanation is that DeepFold focuses on contact patterns near the diagonal of the distance matrix, and ignores contact patterns far from the diagonal (e.g., patch (*i, j*) of the distance matrix shown in Figure 1b). This explanation is also supported by our observations in Section III-B

Employing new attention-based global embedding is another reason why the new ContactLib-ATT model outperforms existing methods. Comparing the results of ContactLib-DNN and ContactLib-ATT in Table II, one can observe that the attention-based global embedding achieves higher accuracies than the simple DNN-based global embedding. Similarly, comparing the results of DeepFold and DeepFold-ATT, the attention-based global embedding outperforms the convolution-based global embedding. These observations are consistent with previous researches showing that attention-based encoder layers yield better results for global feature learning [25].

As shown in Table III, ContactLib-ATT11 is again the most accurate method on the testing dataset. Comparing to the results on the validation dataset, the accuracies of all tested methods are dropped by at least 2.7%. Actually, the accuracy of the TMalign-based [11] nearest neighbor classification (described in Section III-C) is also significantly dropped by at most 11.8%. This suggests that the testing dataset is indeed more challenging than the validation dataset (also discussed in Section III-C), and this could be one explanation for the dropped accuracies. Another explanation is the widely known generalization issue.

**TABLE III.** Accuracies on the testing dataset

|  | Class | Fold | Superfamily |
| --- | --- | --- | --- |
| DeepFold | 87.0±1.6% | 74.4±1.3% | 69.7±0.9% |
| DeepFold-ATT | 90.1±1.2% | 75.8±1.2% | 71.0±1.2% |
| ContactLib-DNN | 88.0±1.3% | 70.6±1.4% | 65.6±1.6% |
| ContactLib-ATT00 | 88.0±1.1% | 74.1±0.9% | 71.0±0.8% |
| ContactLib-ATT01 | 90.2±1.2% | 76.2±1.3% | 73.1±0.9% |
| ContactLib-ATT10 | 89.4±1.3% | 75.4±1.6% | 72.2±1.2% |
| ContactLib-ATT11 | **91.2±0.8%** | **77.0±0.4%** | **74.0±0.6%** |

### B. Understanding when ContactLib-ATT works

To figure out when ContactLib-ATT works, the superfamily classification accuracies on different subsets of the validation dataset are analyzed. In the first experiment, the number of training homologs in the same superfamily of the query protein is counted, and the validation dataset is divided into subsets based on this superfamily size. As shown in Table IV, as the superfamily size increases, the prediction accuracy increases. For the relatively small superfamilies with less than eight members, using H-bond contexts are at least 8.8% more accurate than using full structures. For the relatively large superfamilies with at least eight members, ContactLib-ATT with the attention-based global embedding makes up-to 6.6% more accurate predictions than ContactLib-DNN with the DNN-based global embedding. Therefore, H-bond contexts boost the few-shot learning accuracies, while attention techniques maximize the big-data utilities. This is why ContactLib-ATT outperforms other tested methods despite the superfamily size.

**TABLE IV.** Superfamily classification accuracies for different training superfamily sizes

| Size | 1 | [2, 4) | [4, 8) | [8, 16) | [16, 32) | [32, ∞) |
| --- | --- | --- | --- | --- | --- | --- |
| DeepFold | 21.3% | 43.7% | 59.7% | 69.5% | 82.2% | 88.6% |
| DeepFold-ATT | 19.8% | 42.3% | 60.8% | 76.1% | 86.1% | 90.0% |
| ContactLib-DNN | 26.5% | 49.6% | 62.0% | 75.7% | 80.7% | 85.9% |
| ContactLib-ATT | **30.1%** | **53.8%** | **73.8%** | **83.2%** | **87.9%** | **91.0%** |
| Count | 89 | 173 | 230 | 320 | 442 | 1026 |

In the Second experiment, the validation dataset is divided into subsets based on the query sequence length. As shown in Table V, the prediction accuracy increases as the sequence length increases. Comparing to DeepFold [17], ContactLib-DNN is significantly more accurate for short proteins with less than 64 residues (56.1% v.s. 49.3%) and long proteins with at least 512 residues (88.2% v.s. 80.0%). In fact, our implementation of DeepFold employs six convolution layers with a kernel size of five and a stride step of two. Consequently, each neuron of the last convolution layer covers 253 neurons (i.e., residues) of the input layer. If the query sequence length is significantly smaller than 253, most of the input signals become padding zeros (i.e., noises). If the query sequence length is significantly bigger than 253, remote contact patterns (that are more than 253 residues away on the backbone) cannot be captured by a single neuron. Therefore, the number of convolution layers of DeepFold is a trade-off parameter that cannot optimize both short and long proteins. This problem is well addressed by H-bond contexts that are independent from the depth of the deep learning model.

**TABLE V.** Superfamily classification accuracies for different query sequence lengths

| Length | [20, 64) | [64, 128) | [128, 256) | [256, 512) | [512, ∞) |
| --- | --- | --- | --- | --- | --- |
| DeepFold | 49.3% | 73.8% | 77.1% | 80.3% | 80.0% |
| DeepFold-ATT | 57.3% | 76.8% | 78.4% | 83.1% | 66.0% |
| ContactLib-DNN | 56.1% | 71.2% | 76.5% | 84.6% | **88.2%** |
| ContactLib-ATT | **60.0%** | **80.3%** | **83.7%** | **86.6%** | 87.4% |
| Count | 82 | 740 | 935 | 496 | 27 |

One limitation of the attention-based global embedding is also illustrated in Table V. Similar to nature language processing problems [19], insufficient number (e.g., 27) of long proteins becomes an accuracy bottleneck (87.4% v.s. 88.2%) of attention-based global embedding. This issue might be eased as the number of large protein structures increases with recent developments on Cryo-EM large molecule structure determination techniques [38], [39]. Handling large proteins is also a known challenge for future SCOP releases [1].

### C. Understanding when ContactLib-ATT fails

To figure out why ContactLib-ATT occasionally fails, an interesting case study on SCOP superfamily a.118.8 [1] is discussed in this section. The superfamily is chosen based on the following observations. The superfamily is relatively big with 39, 8 and 10 proteins in the training dataset, the validation dataset and the testing dataset, respectively. Recall that the model is trained five times with random initializations. Using the five trained models, the validation accuracies of the superfamily range between 50% and 88% with an average of 75%. However, the testing accuracies of the superfamily range between 20% and 60% with an average of 32%. In summary, the validation accuracies suggest that there are sufficient training data to train an accurate model, but the testing accuracies are significantly lower.

In order to understand the challenge to classify SCOP superfamily a.118.8, an alignment between a query structure and its nearest (i.e., from the same SCOP family) training homolog is shown in Figure 2. It can be seen that the training homolog is much shorter than the query protein. Actually, eight of the ten query proteins of the testing dataset are more than 100 residues longer than their nearest training homologs. One possible explanation is that these structures contain two domains, which are treated as a single domain in SCOP. Indeed, it is known to be a common error of SCOP [1]. If the SCOP domain partition is correct, this raises new challenges for ContactLib-ATT to make predictions based on fragment instead of global similarities. This problem should be better handled by DeepFold [17] because it primarily depends on fragment similarities.

**Fig. 2.**
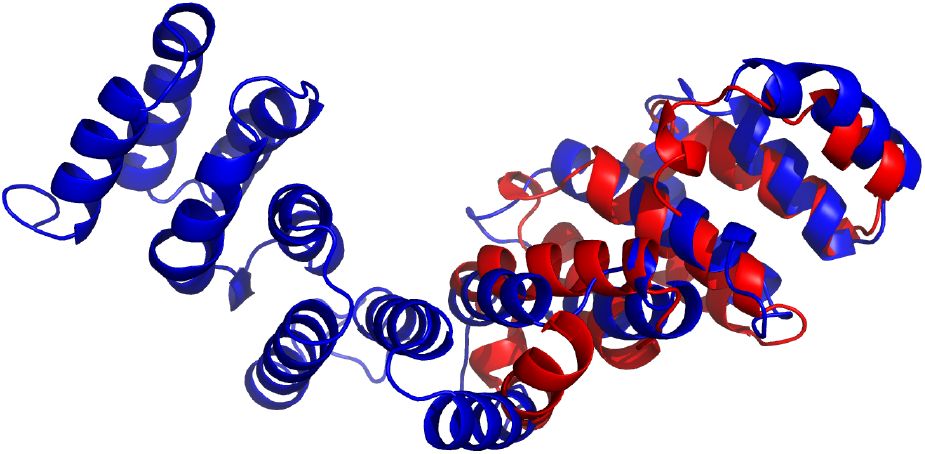
Case study of SCOP superfamily a.118.8: only fragment similarities are observed by TMalign between query protein D2MR3A1 and its nearest training homolog D1RZ4A2, while both proteins belong to SCOP family a.118.8.9.

Consensus approaches can help to ease the fragment similarity problem of SCOP superfamily a.118.8. Our consensus solution is based on the key observation that the true homologs tend to remain in the top-k list if we train and test the model several times, while the false homologs do not share similar trends. Recall that the model is trained five times in our experiments, and each trained model produces a softmax probability distribution. If the average of the five distributions is calculated as the consensus distribution, the testing accuracy is increased from 32% to 60%. If the same consensus method is applied to the experiment shown in Table III, the superfamily classification accuracy will be increased from 74.0% to 76.8%. This suggests us to investigate stochastic weight averaging [40], [41] for ContactLib-ATT, and we would like to investigate more possibilities in future releases.

### D. Homology Search Engine with ContactLib-ATT

In this section, an application of SCOP superfamily classification [1] is analyzed. Specifically, ContactLib-ATT is demonstrated to be a high-quality search engine to find remote homologous proteins within the same SCOP superfamily. To simulate a search engine, ContactLib-ATT and DeepFold [17] is modified to return the top-k candidate list of superfamilies and a randomly selected representative for each candidate superfamily. For references, one sequence-based alignment method (NWalign, [42]) and one structure-based alignment method (TMalign, [11]) is tested. In this experiment, the validation dataset is used as query proteins, and the training dataset is used as the protein database to be searched. As widely accepted, a high-quality search engine should return a candidate list instantly (i.e., in less than a second), and the candidate list should contain at least one true homolog. Thus, the running time and the hit rate (i.e., the probability) that the top-k candidate list includes one true homolog is reported.

As shown in Table VI, if the top-10 list returned by ContactLib-ATT is manually checked by an expert, a protein homolog can be found with a probability of 91.9%. Although different datasets are used, this hit rate is significantly higher than the best results reported by state-of-the-art alignment-free homology search methods [17], [26], [27], [43]. One common feature shared by these alignment-free methods and ContactLib-ATT is that they complete the database search instantly. On the other hand, although structure alignment tools, such as TMalign [11], are more accurate, they are not fast enough as search engines. Moreover, sequence alignment tools, such as NWalign [42], yield low accuracies because sequences are not as reliable as structures to find remote protein homologs. In summary, ContactLib-ATT is the best alignment-free method to implement a search engine for remote protein homologs because it is not only more accurate but also sufficiently fast.

**TABLE VI.** Hit rates and running times as a homology search engine

|  | Top-1 | Top-5 | Top-10 | Top-20 | Time |
| --- | --- | --- | --- | --- | --- |
| NWalign | 0.3% | 2.3% | 5.0% | 9.8% | 3.0 mins |
| TMalign | **90.6%** | **95.6%** | **96.5%** | **97.6%** | 16.3 mins |
| DeepFold | 75.7% | 85.3% | 88.4% | 90.7% | **instant** |
| ContactLib-ATT | 82.4% | 89.5% | 91.9% | 93.8% | **instant** |

## IV Conclusion

In summary, we have introduced ContactLib-ATT, a general-purpose attention-based protein structure embedding framework. When applying on the SCOP superfamily classification problem, ContactLib-ATT achieves an accuracy of 82.4%, which is 6.7% higher than the classic convolution neural network. ContactLib-ATT achieves such significant improvements because it employs a novel two-level embedding approach so that local features and global features are embedded separately. This idea comes from recent Transformer researches on computer vision showing that the best local embedding model is not necessarily the same as the best global embedding model, and attention-based encoder layers should be the first choice for global embedding [25], [44]. Moreover, ContactLib-ATT is an end-to-end deep learning model that does not rely on any database scanning. As a result, classification is done instantly. All these observations well support our conclusion that attention is all you need for general-purpose protein structure embedding.

In the future, we would like explore more possibilities to improve ContactLib-ATT. For example, ContactLib-ATT is capable of handling a variety of input features, such as amino acid features (e.g., residue type), local structure features (e.g., torsion angles), relative structure features (e.g., relative spatial encoding, [22]). Domain partition is also required to handle multi-domain structures properly. Stochastic weight averaging [40], [41] will be implemented, and hopefully better generalization abilities will be demonstrated. We also would like to try to visualize and to understand what is learned by ContactLib-ATT [45]. More applications on protein design, drug design and protein docking will be explored. Finally, we would like to build a structure-based search engine for homologous proteins.

